# Delayed identification of long-overlooked *Aedes* (*Aedimorphus*) *vexans* in Tahiti Island, French Polynesia

**DOI:** 10.1101/2025.08.07.669041

**Authors:** Philippe Boussès, Jérôme Marie, Pierre Kengne, Françoise Mathieu-Daudé, Vincent Manzanilla, Frédéric Lardeux

## Abstract

*Aedes (Aedimorphus) vexans* (Meigen, 1830) is a globally distributed mosquito species commonly associated with temporary flooded habitats, including those in the South Pacific region. While its presence has been documented in several Pacific archipelagos, its occurrence on Tahiti island had remained unconfirmed. A retrospective examination of historical mosquito collections held at the Institut de Recherche pour le Développement (IRD, France) and the Institut Louis Malardé (ILM, Tahiti) revealed evidence of *Ae. vexans* on Tahiti dating back as early as 1993. This historical detection was corroborated by targeted field surveys conducted between 2023 and 2025, which confirmed the species’ presence at multiple sites, particularly in the southern part of the island.

To support these observations, COI gene sequences from newly collected specimens were analyzed alongside reference *Ae. vexans* sequences from GenBank. The Tahitian specimens fall unambiguously within the Pacific clade of *Ae. vexans* and show no genetic divergence that would support their assignment to *Ae. nocturnus* or *Ae. nipponii*. Similarly, sequences published under these names in GenBank do not form distinct lineages within *Ae. vexans*. This study provides the first confirmed record of *Ae. vexans* on Tahiti, based on both morphological identification and molecular evidence. The findings underscore the need to integrate molecular tools into entomological monitoring to improve species detection and prevent misidentifications in island environments where vector diversity is often underestimated.

## Introduction

*Aedes* (*Aedimorphus*) *vexans* (Meigen, 1830) is a widely distributed mosquito species typically associated with floodwater habitats. It is known for its opportunistic biting behaviour and potential involvement in the transmission of various arboviruses (Wilkerson *et al*. 2021). Its ecological plasticity and high dispersal capacity have enabled its successful establishment across a broad spectrum of environments worldwide—from temperate to tropical regions—including several island groups in the Pacific, with the notable exception of South America.

In the South Pacific, *Aedes vexans* has been reported from most archipelagos (Wilkerson *et al*. 2021), but its presence in French Polynesia has remained unconfirmed. A single anecdotal observation from Moorea Island, near Tahiti in the Society Islands, was reported nearly two decades ago (Russell 2004), but subsequent entomological surveys failed to detect the species, leaving its status in the region unresolved (Russell & Burkot 2023). Until now, no formal record existed confirming its presence on Tahiti.

This study reports the first confirmed identification of *Ae. vexans* in Tahiti, based on both diagnostic morphological characteristics and COI barcoding. Despite its global distribution, the presence of *Aedes vexans* in French Polynesia has remained undocumented until this study. Morphological resemblance to other local *Aedes* species, the lack of a regional identification key, and limited sampling efforts in ephemeral or flood-prone habitats may have contributed to this oversight. The study also suggests a long-standing presence of *Aedes vexans* in Tahiti, alongside recent evidence of its local establishment, underscoring the need to re-evaluate the regional diversity of mosquito fauna. Strengthening identification efforts is essential to improve understanding of the species’ ecology and its potential role in arbovirus transmission in French Polynesia.

## Material and methods

### Study area

Tahiti is a volcanic island in French Polynesia, located in the South Pacific Ocean at approximately 17.667°S latitude and 149.500°W longitude. The island features a rugged central volcanic massif, with elevations ranging from sea level to 2 241 m at Mount Orohena, its highest point. A narrow coastal plain, typically 1 to 3 km wide, surrounds the island and consists of gently sloping terrain extending from the shoreline to the base of the volcanic mountains.

Tahiti experiences two distinct seasons. The wet season, from November to April, features temperatures ranging from 25°C to 31°C. Rainfall is significant during this period, with monthly averages between 200 mm and 350 mm, totaling approximately 2500 mm annually. This season is characterized by frequent and intense rainfall, particularly in the central and western regions where the mountainous terrain enhances precipitation. The dry season, from May to October, has average temperatures between 23°C and 29°C. Rainfall decreases substantially during this time, with monthly averages ranging from 50 mm to 150 mm. This season is characterized by lower humidity and reduced precipitation, resulting in generally drier conditions.

Tahiti’s terrestrial ecosystems are diverse, featuring lush rainforests, rugged volcanic mountains, and ecologically important coastal wetlands. The island’s altitudinal gradient supports a range of vegetation types, from lowland forests to montane and cloud forests. In contrast, human-modified landscapes—primarily along the coast—reflect the influence of urban development and agricultural activities.

### Field collections of *Ae. vexans*

Pre-imaginal stages of *Ae. vexans* were collected during entomological surveys conducted in April–May 2023, April and August 2024, and February 2025, across four municipalities along the southeastern coast of Tahiti—Paea, Papara, Mataiea, and Atimaono—covering a stretch of approximately 20 km of coastline. Larval sampling targeted shallow temporary and permanent ponds, including sunlit or shaded grassy depressions subject to flooding, which represent the preferred habitats of this species (Russell & Burkot 2023). Multiple mosquito species were recorded during these surveys. For each sampling event, fourth-instar (L4) larvae were sorted by species and either preserved in 70% ethanol for subsequent molecular analysis or prepared as slide-mounted specimens for morphological study. Remaining larvae and pupae were reared to the adult stage and preserved as pinned specimens for entomological collections.

### Search for *Ae. vexans* in laboratory collections

*Aedes vexans* was also sought among historical specimens housed in two entomological collections containing material from Tahiti: (i) the collection of the Institut Louis Malardé (ILM) in Tahiti and (ii) the ARIM (Arthropodes d’Intérêt Médical) collection at IRD-France (ARIM 2024). This retrospective examination considered the possibility that *Ae. vexans* might have previously been misidentified or confused with other species, such as *Aedes edgari* (Stone & Rosen, 1952). Consequently, all voucher specimens were thoroughly examined.

### Morphological identification of *Ae. vexans*

Field-collected larvae, emerged adult specimens and collection vouchers were morphologically identified using dichotomous keys specific to the Pacific region (Bohart & Ingram 1946; Rageau 1958; Belkin 1962; Huang 1977), taxonomic group revisions, and original species descriptions (Stone & Rosen 1952; Belkin 1962). Adult specimens of *Ae. vexans* and *Ae. edgari* exhibit relatively similar morphological features, which may lead to misidentification. To minimize this risk, key diagnostic characters were carefully examined across the larval, adult female and adult male stages, as detailed below:

In the larval stage of *Ae. vexans*, the comb on abdominal segment VIII consists of 6–12 spines that are imperfectly aligned, with the two terminal pecten teeth distinctly separated from the others. Setae 1-S are short, inserted beyond the pecten, and the siphon is relatively long. In contrast, *Ae. edgari* exhibits a comb on segment VIII composed of a patch of numerous scales, regularly spaced pecten teeth, well-developed setae 1-S inserted at the apex of the pecten, and a significantly shorter siphon (Fig. 2).

**Figure 1.**
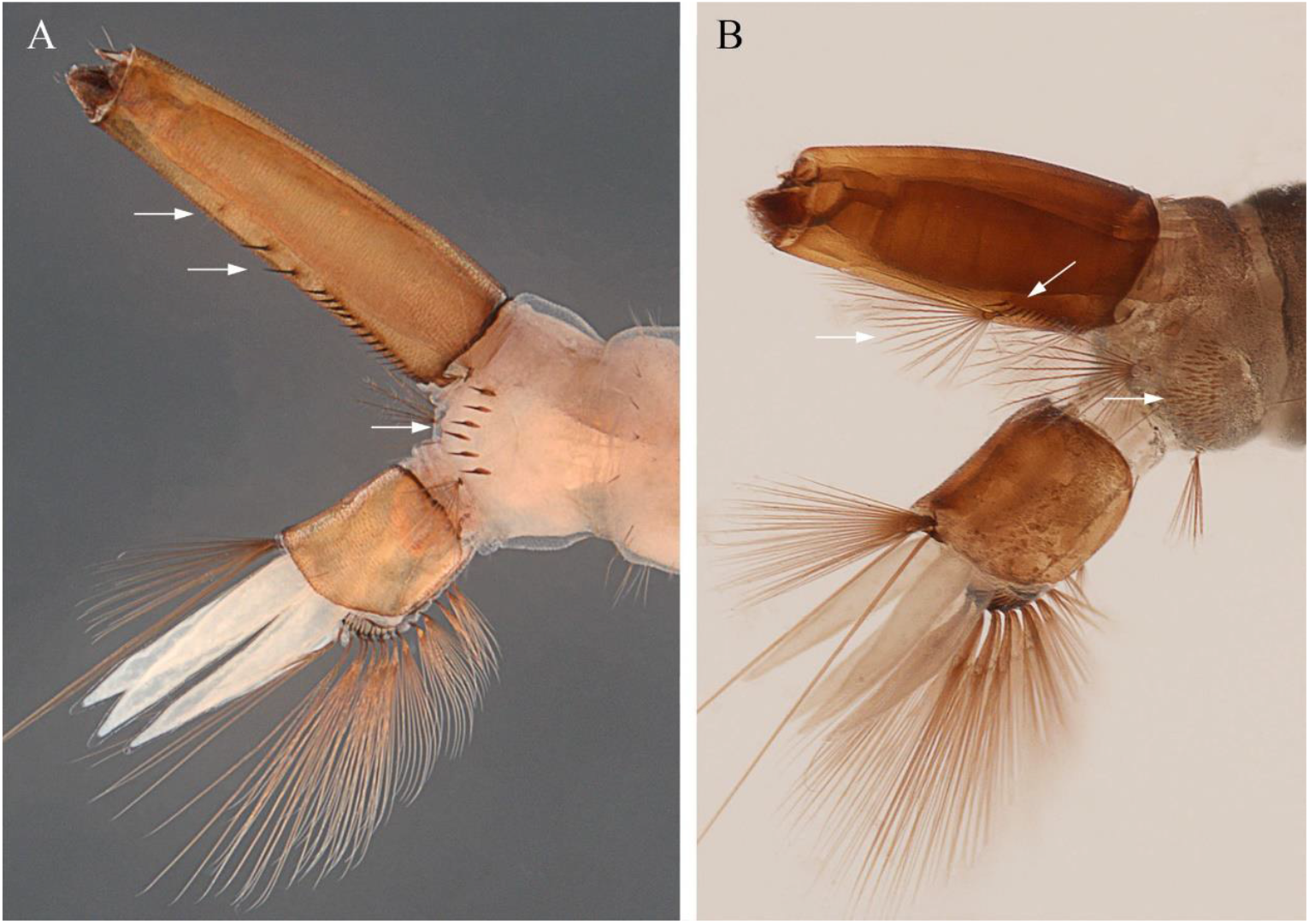
Morphological characteristics used to distinguish between larvae of *Ae. vexans* (A) and *Ae. edgari* (B). Specimens are from sample PhB-2023-012 of Papara. Segment VIII-X: comb VIII formed of 6-12 more or less aligned spines, the two terminal teeth of the pecten clearly separated from the others, setae 1-S short inserted beyond the pecten, relatively long siphon in *Ae. vexans* (A), and comb VIII made up of a patch of scales, pecten teeth regularly spaced, well-developed 1-S setae inserted at the apex of the pecten and siphon significantly shorter in *Ae. edgari* (B).

**Figure 2.**
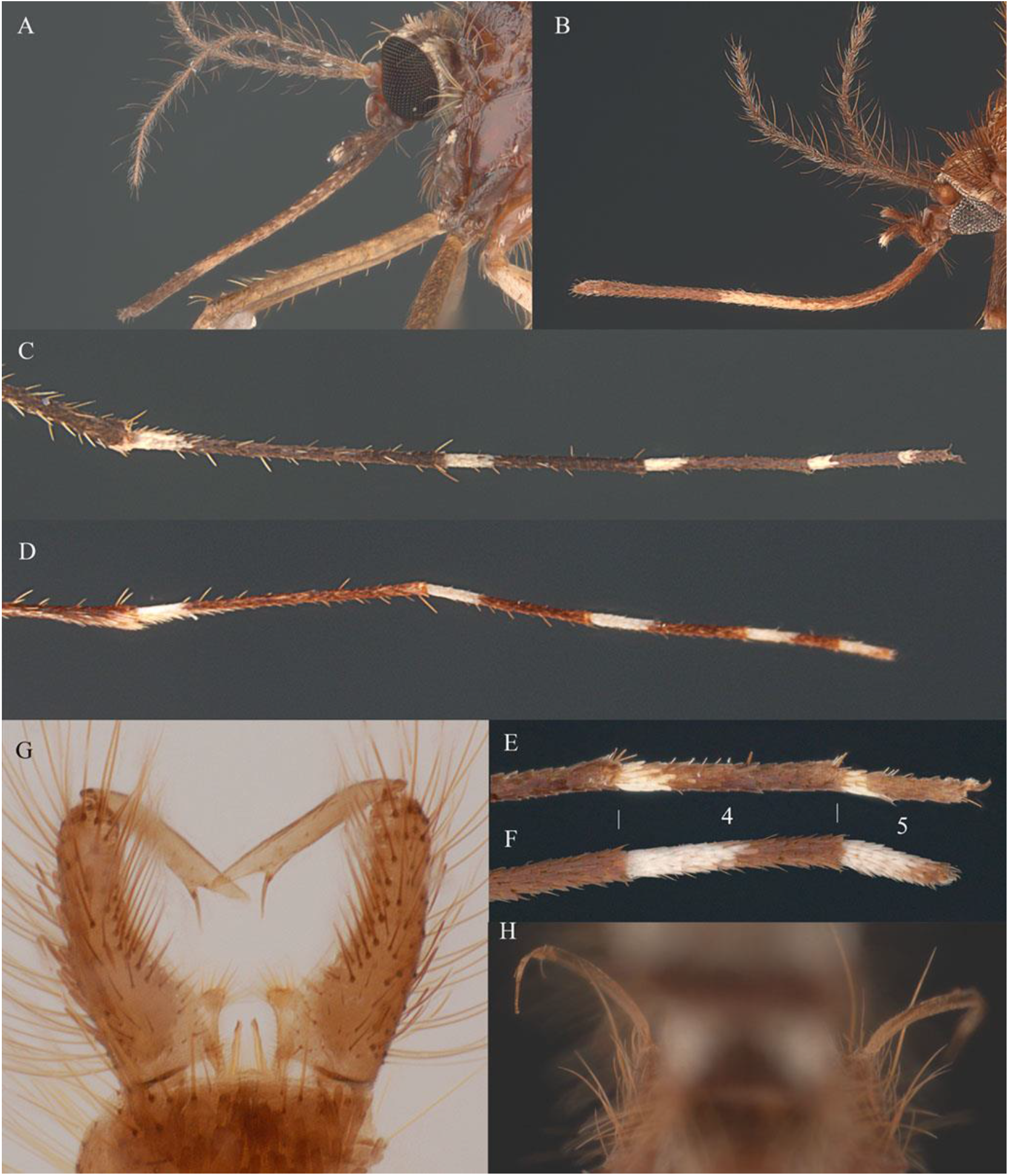
Morphological characteristics used to distinguish between adults of *Ae. vexans* and *Ae. edgari*. Female. Ornamentation of the proboscis: with a very wide incomplete median pale band not sharply defined in *Ae. vexans (*A), and with a large clear median third pale ring in *Ae. edgari* (B). Ornamentation of tarsomere 5 (III): pale scales restricted to the 1/3 of the basal area in *Ae. vexans* (C, E), and almost entirely covered with pale scales, with just some dark scales at the apex, in *Ae. edgari* (D, F). Male genitalia. Form of gonostylus: gradually expanded to apex. Spine of gonostylus articulated subapically, straight in *Ae. vexans* (G), and tapered gonostyle thinning from the base to the apex and extended apically by a sharp spine in *Ae. edgari* (H).

In females, the proboscis ornamentation of *Ae. vexans* features a wide but incomplete median pale band that is poorly defined. In contrast, *Ae. edgari* exhibits a large, distinct pale ring located in the median third of the proboscis. In *Ae. vexans*, ornamentation of tarsomere 5 of the hind leg (5-III) is restricted to pale scales on the basal third, whereas in *Ae. edgari*, this segment is almost entirely pale-scaled except for a dark apical area.

In males, the gonostylus of *Ae. vexans* gradually expands toward the apex, with a straight spine articulated subapically. In contrast, the gonostylus of *Ae. edgari* tapers from the base to the apex and ends in a sharp apical spine (Fig. 2).

### Molecular identification

Genomic DNA was extracted from four field-collected larvae originating from samples PhB-2023-010 and PhB-2023-012 (Tab. 1), using the DNeasy Blood and Tissue Kit (Qiagen, Germany), according to the manufacturer’s protocol. DNA pellets were air-dried and resuspended in 50 µL of nuclease-free water. The mitochondrial cytochrome c oxidase subunit I (COI) gene was amplified using the universal primers LCO1490 (forward: 5′-GGT CAA CAA ATC ATA AAG ATA TTG G-3′) and HC02198 (reverse: 5′-TAA ACT TCA GGG TGA CCA AAA AAT CA-3′) (Folmer *et al*. 1994).

PCR reactions were carried out in a final volume of 25 µL, containing 1.5 mM MgCl_2_, 0.2 mM of each dNTP (Eurogentec), 10 pmol of each primer, 1× Taq Polymerase buffer, 1 U of Taq DNA Polymerase (Qiagen, Courtaboeuf, France), and 1–5 ng/µL of template DNA. Thermal cycling conditions included an initial denaturation at 95 °C for 3 min, followed by 35 cycles of denaturation at 94 °C for 30 s, annealing at 50 °C for 30 s, and extension at 72 °C for 2 min, with a final extension step at 72 °C for 10 min.

PCR amplicons were sequenced bidirectionally using Sanger sequencing on an ABI 3130XL Genetic Analyzer (Eurofins Genomics, France). Resulting chromatograms were manually inspected and edited to correct sequencing errors. The sequences were then assembled and aligned using MUSCLE (Edgar 2004). All four Tahitian COI sequences of 676 bp were identical, representing a single haplotype. For preliminary taxonomic identification, the sequence was queried against the NCBI GenBank database using the BLAST algorithm. Sequences with 100% query cover and 100% identity were *Ae. vexans*. Other sequences were retrieved with >95% query coverage and >93% sequence identity, yielding over 400 matches. All retrieved sequences were aligned and trimmed; those exhibiting ambiguous bases or incomplete coverage were excluded, resulting in a curated dataset of 389 sequences. To reduce redundancy and simplify phylogenetic analysis, VSEARCH clustering (Rognes *et al*. 2016) was applied with a centroid identity threshold of 0.990, producing 47 representative haplotype groups. Most haploids were assigned to *Aedes vexans* though some lacked precise taxonomic identification but were retained to preserve genetic diversity. Several sequences of *Aedes hirsutus* (Theobald, 1901), a closely related species within the subgenus *Aedimorphus*, were included as an outgroup. The four Tahitian sequences, three *Ae. vexans* sequences from New Caledonia, and one sequence from Moorea Island—collected in December 2006 and initially identified only as *Culicidae* sp. sc_01371—(Ramage *et al*. 2017) but sharing 100% identity and coverage with the Tahitian sequences were manually added to this dataset. These Pacific Island sequences were excluded from VSEARCH clustering to maintain intraspecific diversity within this geographic group. The final dataset was analyzed in MEGA 7 (Kumar *et al*. 2016), where the Tamura 3-parameter model with gamma distribution (T92+G) (Tamura 1992), was identified as the best-fit substitution model. A Maximum Likelihood phylogenetic tree was then constructed with 500 bootstrap replicates to evaluate node support (Felsensten 1985).

### Data preservation and storage

All field samples—including collected larvae, field-caught adults, and reared adults (emerged from field-collected larvae)—have been deposited in the entomological collection of the Institut Louis Malardé, Tahiti, French Polynesia. The four *Ae. vexans* COI sequences from Tahiti were deposited in GenBank under accession numbers PV805667.1 and PV807927.1 for the two specimens of the PHB-2023-010 sample, and PV807926.1 and PV807925.1 for the two specimens of the PHB-2023-012 sample.

## Results

### Field samples

In 2023, *Ae. vexans* was collected from larval habitats located near the main roadside at sea level altitude in Mataeia, Papara, and Atimaono (Table 1). In Mataeia, larvae were found on 26.IV.2023 in various shallow depressions within a *Lolium perenne* L. lawn adjacent to a sun-exposed football field. *Aedes vexans* was the only species present, with abundant larvae predominantly at the L4 developmental stage and a few pupae. A subsequent visit to the site on 27.V.2023 yielded no larvae. In Papara, *Ae. vexans* larvae were collected on 27.IV.2023 from a small shallow, mostly shaded pond with dense vegetation of *Commelina diffusa* N.L. Burman. Numerous *Ae. vexans* L4 larvae were present, along with a single captured *Ae. edgari* larva. However, 12 days later (09.V.2023), neither species was found at the site; instead, the habitat was dominated by *Culex annulirostris* Skuse, 1889, and *Culex quinquefasciatus* Say, 1823, both present in high numbers. In Atimaono, no larval habitats positive for *Ae. vexans* were detected. However, two adult females were opportunistically collected around midday while attempting to bite during ongoing field surveys.

The following year, on 09.VIII.2024, a new *Ae. vexans* larval site was discovered in Paea. The habitat consisted of a sunlit grassy area planted with a mix of *Alternanthera sessilis* (Linnaeus) R. Brown ex A.P. de Candolle 1813, *Paspalum paniculatum* Linnaeus, and *P. conjugatum* P. J. Bergius 1762. A water leak had created a large, shallow puddle in a ground depression, where *Ae. vexans* was the sole mosquito species present. Larvae were abundant, predominantly at the L4 developmental stage, with a few pupae. On 22.IV.2024, numerous *Ae. vexans* larvae were again found in the grassy depressions at the Mataeia site, all at the L2 developmental stage. Additionally, a new larval site was identified within the same municipality. This newly discovered habitat, located in a temporary puddle formed in a grassy depression planted with a mix of *Paspalum* sp. and young *A. sessilis*, was situated away from the public beach (Figure 3). The site was notable for its abundance of *Ae. vexans* larvae, predominantly at the L4 stage.

**Table 1.** Chronological list of mosquito collections in Tahiti, including one sample in which *Aedes vexans* was not detected but is discussed in the text. Sample ID (when assigned), Municipality where the sample was collected, GPS data in decimal degrees, collection date, collector name, habitat type and collected species. Asterisk indicates samples from which larvae were used for mitochondrial cytochrome c oxidase subunit I (COI) gene sequencing (two larvae per sample).

| Sample ID | Municipality | GPS data<br>(lat., long.) | Collection<br>date | Collector | Habitat | Collected species |
| --- | --- | --- | --- | --- | --- | --- |
| FLX-1993-001<br>In ARIM collection | Paea | - | 1993 | Lardeux F. | Inundated grassy area | *Ae. vexans* |
| In ILM collection | Papara | - | V.2000 | Séchan Y. | Unknow | *Ae. vexans* |
| PhB-2023-010*<br>In ILM collection | Mataiea | 17.76478 S<br>149.41023 W | 26.IV.2023 | Boussès P., Marie J. | Small depressions within a lawn (sports field) | *Ae. vexans* |
| PhB-2023-012*<br>In ILM collection | Papara | 17.75768 S<br>149.51211 W | 27.IV.2023 | Boussès P., Marie J. | Shallow, shaded pond with dense vegetation | *Ae. vexans*<br>*Ae. edgari* |
| PhB-2023-015<br>In ILM collection | Papara | 17.75768 S<br>149.51211 W | 09.V.2023 | Boussès P., Marie J. | Shallow, shaded pond with dense vegetation<br>(same site as PhB-2023-012) | *Cx. annulirostris*<br>*Cx. quinquefasciatus* |
| In ILM collection | Atimaono | 17.77887 S<br>149.45874 W | 23.V.2023 | Mathieu-Daudé F. | Biting female | *Ae. vexans* |
| In ILM collection | Paea | 17.73249 S<br>149.57828 W | 09.VIII.2024 | Mathieu-Daudé F. | Sunlit grassy area with a large, shallow<br>puddle | *Ae. vexans* |
| In ILM collection | Mataiea | 17.77363 S<br>149.41158 W | 22.IV.2024 | Mathieu-Daudé F. | Grassy area with shallow puddles near beach | *Ae. vexans* |
| In ILM collection | Paea | 17.72896 S<br>149.58046 W | 19.II.2025 | Marie J. | Grassy area with shallow puddles | *Ae. vexans* |
| In ILM collection | Paea | 17.72925 S<br>149.58021 W | 19.II.2025 | Marie J. | Grassy area with shallow puddles with<br>decomposing leaf litter | *Ae. vexans*<br>*Ae. edgari* |

**Figure 3.**
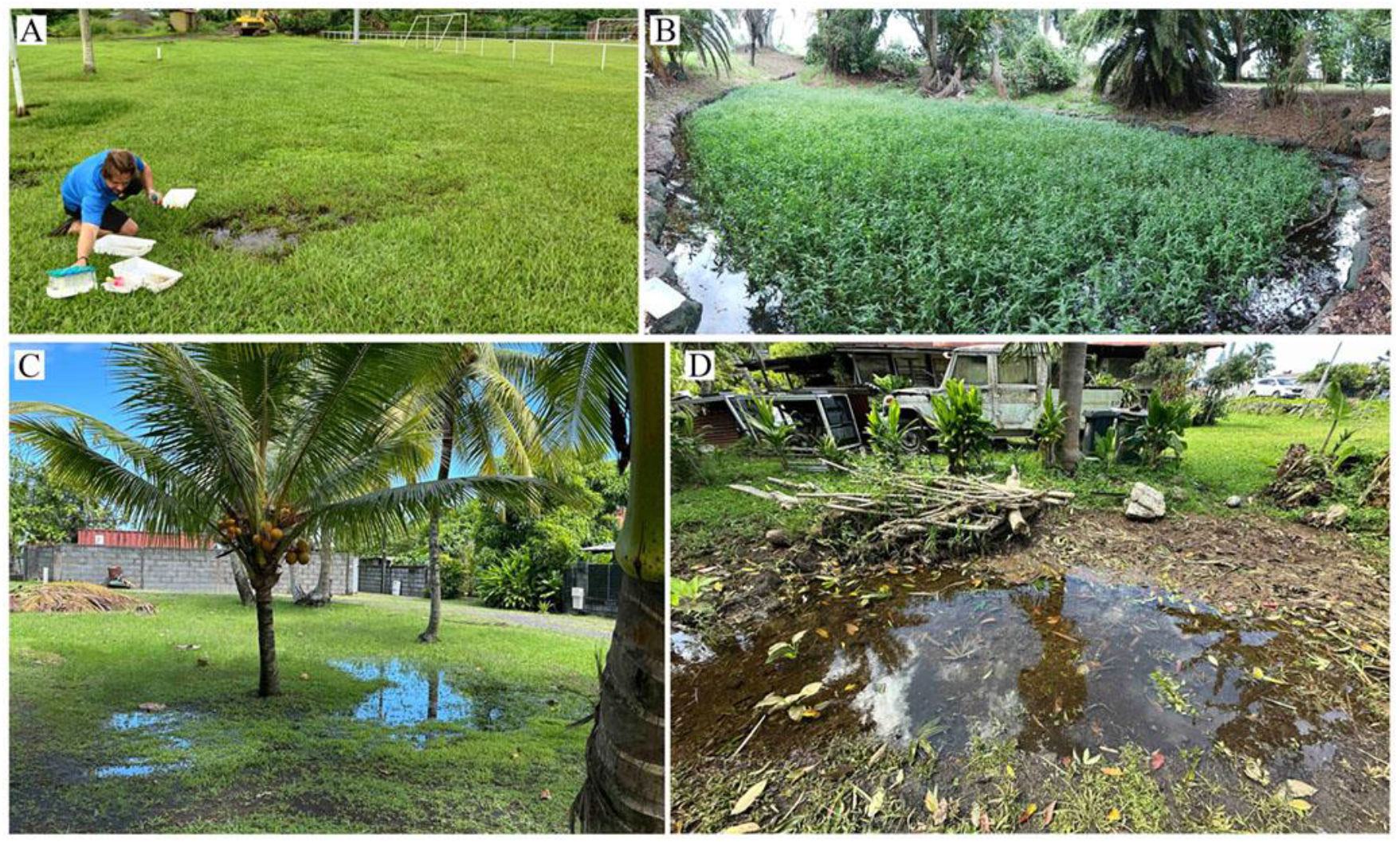
Larval habitats of collected *Ae. vexans* during this study. (A) Mataeia site (2023) (PHB-2023-010), (B) Papara site (2023) (PHB-2023-012), (C) Mataeia site (2024), and (D) Paea site (2025). In site B and D, *Ae. vexans* was collected together with *Ae. edgari* (photo A includes one of the authors of this article).

In February 2025, following a period of heavy rainfall, two temporary *Aedes vexans* breeding sites were identified in close proximity within the commune of Paea. Both consisted of shallow, grass-covered ground depressions that retained standing water for 5 to 6 days. At one of these sites, *Ae. edgari* was also observed developing, though at a much lower density than *Ae. vexans*. The puddle, approximately 5–6 cm deep and 2 meters in diameter, reached a maximum water temperature of 35 °C and contained herbaceous vegetation as well as decomposing leaf litter (Figure 3).

### Specimens from laboratory collections

The oldest known sample of *Ae. vexans* in Tahiti is a set of eight stage IV larvae collected by F. Lardeux in 1993 (day and month not recorded on the sample vial) from a flooded grassy area near the main roadside at sea level altitude in Paea. Preserved in 70% alcohol and initially misidentified as *Ae. edgari*, the sample—stored in the ARIM collection at IRD-France—was later morphologically confirmed to be *Ae. vexans*. In this collection, which includes 498 voucher specimens from French Polynesia, including 262 labeled from Tahiti (113 preimaginal and 149 adults), collected on various occasions by J. Rageau (1959), J.M. Klein (1981–1983), J. Brunhes (March 2002), and other unnamed collectors, no additional *Ae. vexans* specimens were identified.

Reviewing the collection at the Entomology Laboratory of the Institut Malardé, Tahiti, revealed a small number of specimens. A set of six slides labeled ‘*Aedes edgari, Papara, May 2000, Y. Séchan coll*.’ were identified, upon examination, as *Ae. vexans* larvae.

### Molecular identification of *Ae. vexans*

A Maximum Likelihood phylogenetic tree is presented in Figure 4 in which bootstrap values below 0.70 are not displayed, and the countries of origin for all specimens corresponding to each haplotype are indicated. Phylogenetic analysis based on COI sequences confirms that the four specimens from Tahiti belong to the *Ae. vexans* species. These specimens, which share an identical haplotype, form a tight monophyletic cluster with sequences from Moorea and New Caledonia within a well-supported Pacific group (bootstrap = 80%). This Pacific group shows no indication of species-level divergence from other *Ae. vexans* lineages and is genetically distinct from *Ae. hirsutus*, which forms a separate, strongly supported clade (bootstrap = 99%), thereby confirming its species status. The Pacific group is closely related to several *Ae. vexans* sequences from East Asia, particularly individuals designated as *Ae. vexans nipponii* (Theobald, 1907) from Japan. Together, they form a moderately supported grouping that also includes sequences from Thailand, South Korea and China. This structure suggests that Pacific populations are more closely related to East Asian lineages than to European or North American ones, pointing toward an eastern origin or dispersal pathway for the Pacific populations.

**Figure 4.**
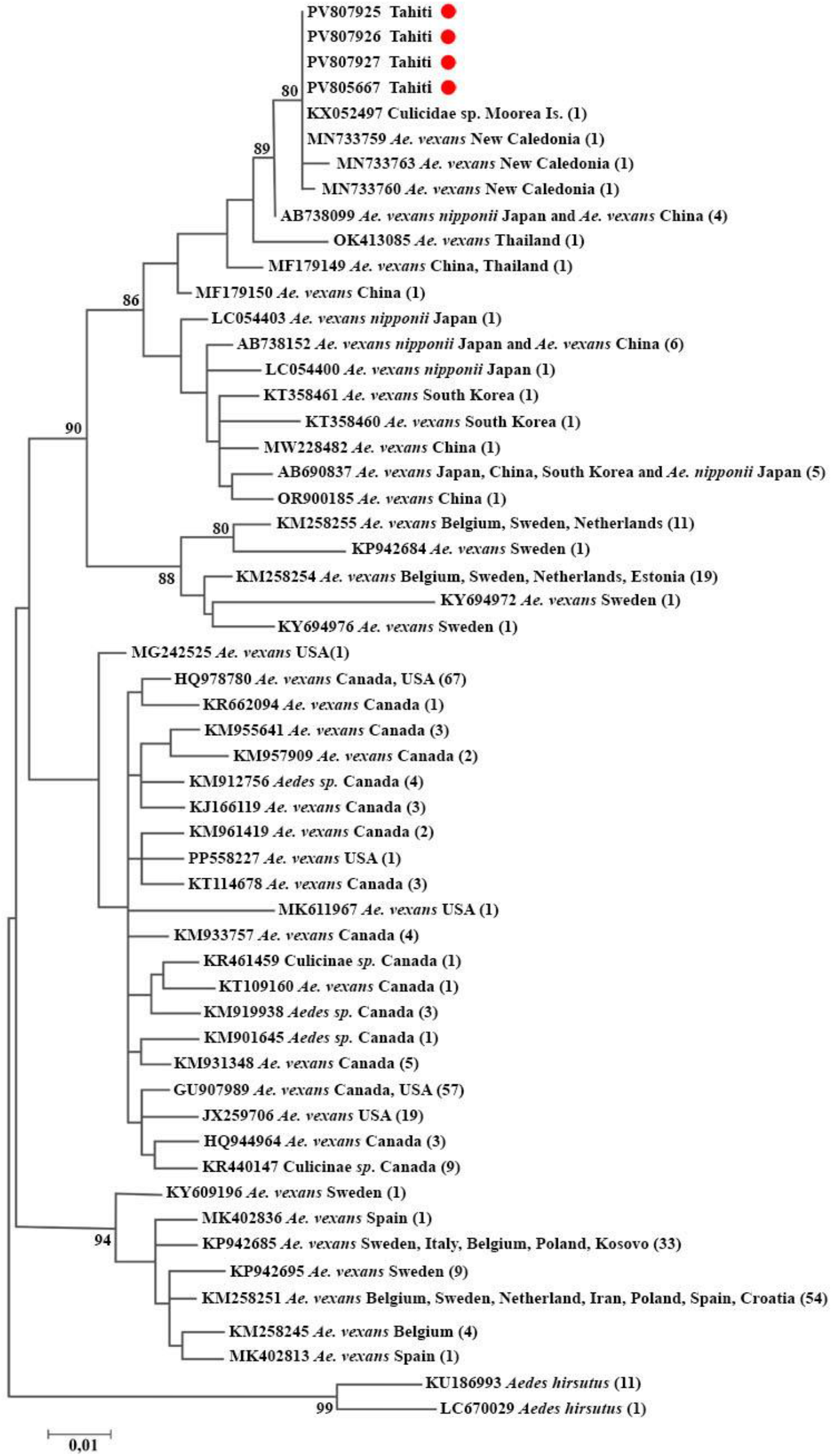
Molecular Phylogenetic analysis by Maximum Likelihood method. The evolutionary history was inferred by using the Maximum Likelihood method based on the Tamura 3-parameter model (Tamura, 1992)The tree with the highest log likelihood (−2333.40) is shown. The percentage of trees in which the associated taxa clustered together is shown next to the branches. Only values >0.70 are shown. Initial tree(s) for the heuristic search were obtained automatically by applying Neighbor-Join and BioNJ algorithms to a matrix of pairwise distances estimated using the Maximum Composite Likelihood (MCL) approach, and then selecting the topology with superior log likelihood value. A discrete Gamma distribution was used to model evolutionary rate differences among sites (5 categories (+*G*, parameter = 0.1766)). Branch lengths are measured in the number of substitutions per site. The analysis involved 55 nucleotide sequences. There was a total of 587 positions in the final dataset The Tahitian sequences are indicated by red dots.

Consistent with the geographic structuring reported by Vargas-Espinosa & Aguirre-Obando (2022), the *Ae. vexans* sequences analyzed in this study cluster according to broad regional origins. Apart from the Asian group described above, European sequences form two distinct and strongly supported groups. One includes samples from Belgium, Sweden, the Netherlands and Estonia (bootstrap = 90%), while the other clusters additional Belgian and Swedish sequences with those from Spain, Italy, Poland, Kosovo, Iran, Netherlands and Croatia (bootstrap = 94%). North American sequences (from Canada and the USA) also form a geographically coherent, though moderately supported, assemblage, reflecting regional clustering with some internal genetic heterogeneity.

In contrast, *Ae. vexans nipponii* sequences are paraphyletic and scattered across multiple subgroups within the “Asian” group, some clustering with Pacific Island samples (e.g., AB738099), others with East Asian sequences from Japan, China, or South Korea. The absence of a monophyletic *nipponii* lineage and the weak internal bootstrap support suggest that this designation reflects geographic labeling or intraspecific variation rather than a valid taxonomic entity.

## Discussion

*Aedes vexans* had long been absent from the Culicidae species list for French Polynesia until now (Rivière 1988). This absence can be attributed to the species not being recognized as present in French Polynesia, as reflected in Belkin’s seminal 1962 reference book (Belkin 1962), which long served as a foundational resource for Tahitian entomologists. This oversight led to the exclusion of *Ae. vexans* from local identification keys (Rivière *et al*. 1979) and resulted in its systematic misidentification as *Ae. edgari*, as occurred with specimens collected in Tahiti in 1993 by Lardeux and in 2000 by Séchan. Indeed, the local key available at the time distinguished only among three recognized *Aedes* species - *Aedes aegypti* (Linnaeus, 1762*), Ae. polynesiensis* Marks, 1951, and *Ae. edgari* - using characters that did not discriminate *Ae. vexans*. Consequently, the key inevitably and unambiguously led to the misidentification of *Ae. vexans* as *Ae. edgari*. Moreover, entomological research in Tahiti has, for decades, primarily focused on vector studies—particularly *Ae. polynesiensis* and *Ae. aegypti*—rather than on mosquito taxonomy. This lack of specific attention to taxonomy has contributed to the oversight in identifying the full diversity of the local Culicidae fauna.

The first published record of the *Ae. vexans* in French Polynesia dates back to November 2003, when two females were collected on the neighboring island of Moorea approximately 17 km northwest of Tahiti, using a light trap (Russell 2004). Unfortunately, these specimens were not preserved (Russell, pers. comm.), and without voucher specimens, the information could not be verified. Therefore, until now, the presence of *Ae. vexans* has never been conclusively confirmed in the Society Islands, leaving uncertainty about whether the species is truly established in the region (Russell & Burkot 2023). Consequently, published Culicidae lists for French Polynesia since the 2003 discovery in Moorea either acknowledge the presence of the species without further discussion (Marie & Bossin 2013; Lecollinet *et al*. 2022) or exclude it altogether (Russell & Burkot 2023).

The present study reports the presence of *Ae vexans* on Tahiti and demonstrates that it has been established there since at least 1993, when it was first collected. However, the date of colonization of Tahiti by this species remains unknown. The 1993 record likely represents a conservative estimate, especially considering the long-standing presence of *Ae. vexans* on numerous Pacific islands. American entomologists had already identified the species in the region as early as the 1930s, and subsequent surveys throughout the mid-20th century confirmed its presence in most archipelagos (Lee *et al*. 1982), including the nearest archipelago, the Cook Islands, approximately 1150 km west of Tahiti (McKenzie 1925; Belkin 1962). These surveys were part of broader entomological efforts, often conducted in connection with disease control programs, such as the American-led campaigns against lymphatic filariasis in the 1940s and 1950s in Tahiti. In this context, it is particularly striking that *Ae. vexans* was not reported from Tahiti during that period, despite intensive mosquito investigations. Notably, *Ae. edgari* was rediscovered and described during the same period. This absence raises the question: was *Ae. vexans* truly absent from Tahiti in the mid-20th century, or was it simply overlooked? *Aedes vexans* is not included in any Culicidae lists for Tahiti up to 2004, and is also absent from the limited voucher material of the Rageau collection from 1959 and the Klein collection from 1981-83, both housed at ARIM-France. However, *Ae. edgari*, which can share similar larval habitats with *Ae. vexans*, is present in both collections. This raises the question once again: was *Ae. vexans* truly absent from Tahiti in the 1980s? A definitive answer cannot be provided.

Recent field sampling in 2023 and 2025 confirms that the species is now well established on Tahiti and occurs across multiple locations on the island. When detected, the species can occur at high larval densities in suitable habitats, although its presence appears limited to brief temporal windows likely driven by short-lived environmental conditions. However, these recent samplings were limited to the southeastern coastal plain of the island. Therefore, while it is challenging to generalize the species’ presence across the entire island, this hypothesis is highly plausible. The distribution of *Ae. vexans* spans the Australasian region and the islands of the western and southern Pacific, extending as far east as the Cook Islands and including Hawaii (Russell & Burkot 2023), where it was introduced accidentally via air transport in January 1962 (Joyce & Nakagawa 1963). Moorea and Tahiti should now be formally added to this geographic distribution list.

Notably, at each collection site in Tahiti, *Ae. vexans* larvae were almost all at the same developmental stage, and subsequent visits did not detect the species again. These observations are consistent with the species’ described bioecology, which, like *Ae. edgari*, breeds in temporary ground pools. The species lays eggs that can remain viable for months on grass or soil and hatch synchronously when flooding occurs, resulting in sporadic but abundant larval populations (Joyce & Nakagawa 1963).

The identification of *Ae. vexans* in Tahiti raised the taxonomic question of whether these populations should instead be referred to as *Ae. nocturnus* (Theobald, 1903), a name historically applied to similar Pacific forms. Some authors have treated *Ae. nocturnus* as a synonym or subspecies of *Ae. vexans* (Reinert 1973; Wilkerson *et al*. 2021), while others have considered it a distinct species based on its geographical occurrence across the Australasian and Pacific regions (Lee *et al*. 1982; Harbach & Wilkerson 2023). However, previous morphological criteria used to distinguish *Ae. nocturnus*, particularly at the larval stage, have been shown to be unreliable (Reinert 1973), and, to date, no molecular data have supported its separation as a distinct taxon. In our study, COI sequence analysis places Tahitian specimens within a well-supported group of Pacific *Ae. vexans*, with no evidence of species level divergence. These results do not support the recognition of *Ae. nocturnus* as a valid taxon. Consequently, we assign the Tahitian specimens to *Ae. vexans* and consider the use of the name *nocturnus* for these populations to be taxonomically unwarranted. Similarly, *Ae. vexans nipponii*, currently treated as a distinct species (*Aedes nipponii*) by Harbach & Wilkerson (2023) due to minor morphological differences for each stage, does not appear to constitute a coherent or genetically distinct taxonomic unit.

The low COI genetic divergence observed among *Ae. vexans* specimens from Tahiti, New Caledonia, and Japan, suggests a close phylogenetic relationship despite the considerable geographic distances separating these locations. These nearly identical haplotypes, forming a distinct, well-supported group within the broader *Ae. vexans* clade, are consistent with a recent colonization scenario involving a common East Asian source population. The presence of such genetically similar individuals across isolated Pacific islands points to human-mediated dispersal as the most plausible explanation, likely through maritime or aerial transport of immature stages or adult mosquitoes. This interpretation is further supported by the ecological plasticity of *Ae. vexans* and its association with anthropogenic habitats. While long-distance natural dispersal cannot be entirely excluded, the observed genetic uniformity and distribution pattern argue strongly in favor of recent, human-facilitated introductions, reflecting historical and contemporary trade connections across the Pacific.

The low genetic variability among the “Pacific” sequences may also reflect a founder effect, repeated introductions from the same source population, or ongoing gene flow among these regions. It is also important to consider that the COI marker may lack the resolution needed to detect subtle population structure among recently established insular populations. Further investigation using nuclear markers or whole-genome data across a broader geographic sampling range would be necessary to refine the phylogeographic history of *Ae. vexans* in the Pacific region.

## Conclusion

This study provides the first confirmed identification of *Aedes (Aedimorphus) vexans* in Tahiti, based on a combination of morphological characters and COI barcoding. Although previously unrecognized in the local fauna, retrospective examination of entomological collections revealed specimens dating back to 1993, suggesting that the species has likely been established on the island for at least three decades. Field surveys conducted between 2023 and 2025 confirm its current distribution, particularly in the southern part of the island, where it appears to be well established. Phylogenetic analysis places the Tahitian specimens within a well-supported Pacific group of *Ae. vexans*, providing no evidence to support the recognition of *Aedes nocturnus* as a distinct species. The delayed detection of *Ae. vexans* highlights the need to broaden the scope of entomological surveillance in French Polynesia. Although regular weekly monitoring is conducted by public health services across numerous communes, it has traditionally focused on confirmed vectors such as *Ae. aegypti* and *Ae. polynesiensis*. As a result, other *Aedes* and *Culex* species may remain under-detected. Continued field surveillance, combined with modern molecular identification techniques, will be essential to refine the regional inventory of Culicidae species and to evaluate the potential epidemiological implications of newly detected or previously overlooked taxa.

## Acknowledgments

We thank Philippe Marmey (UMR EIO, IRD – Université de la Polynésie française) for his assistance in identifying plant species associated with *Ae. vexans* larval habitats, and Christian Barnabé (UMR INTERTRYP, IRD-CIRAD-UM-II) for his valuable help with COI sequence analysis and phylogenetic reconstruction.

